# Gene expression data support the hypothesis that *Isoetes* rootlets are true roots and not modified leaves

**DOI:** 10.1101/2019.12.16.878298

**Authors:** Alexander J. Hetherington, David M. Emms, Steven Kelly, Liam Dolan

## Abstract

Rhizomorphic lycopsids are the land plant group that includes the first giant trees to grow on Earth and extant species in the genus *Isoetes*. Two mutually exclusive hypotheses account for the evolution of terminal rooting axes called rootlets among the rhizomorphic lycopsids. One hypothesis states that rootlets are true roots, like roots in other lycopsids. The other states that rootlets are modified leaves. Here we test predictions of each hypothesis by investigating gene expression in the leaves and rootlets of *Isoetes echinospora*. We assembled the *de-novo* transcriptome of axenically cultured *I. echinospora*. Gene expression signatures of *I. echinospora* rootlets and leaves were different. Furthermore, gene expression signatures of *I. echinospora* rootlets were similar to gene expression signatures of true roots of *Selaginella moellendorffii* and *Arabidopsis thaliana*. RSL genes which positively regulate cell differentiation in roots were either exclusively or preferentially expressed in the *I. echinospora* rootlets, S. *moellendorffii* roots and *A. thaliana* roots compared to the leaves of each respective species. Taken together, gene expression data from the *de-novo* transcriptome of *I. echinospora* are consistent with the hypothesis that *Isoetes* rootlets are true roots and not modified leaves.

## Introduction

The first giant (> 50 m) trees to grow on Earth, the arborescent clubmosses, were tethered to the ground by rooting structures termed stigmarian systems whose homology has been debated for more than 150 years^1–9^. Stigmarian rooting systems consisted of two components, a central axis (rhizomorph) on which developed large numbers of fine axes (rootlets). There are two competing hypotheses to explain the origin of stigmarian rootlets which we designate, the lycopsid root hypothesis and the modified shoot hypothesis. The lycopsid root hypothesis posits that rootlets are homologous to roots of other lycopsids. The modified shoot hypothesis posits that rootlets are modified leaves (microphylls) and homologous to the leaves of other lycosids.

Stigmarian rootlets were interpreted as true roots by the majority of authors until the mid 20^th^ century^5,6,10–14^. However, a suite of fossil findings in the second half of the 20^th^ century, including fossil embryos, rhizomorph apices and the abscission of rootlets^3,4,15–19^ led to the revival of the modified shoot hypothesis first suggested in 1872, which interpreted rootlets as modified leaves^7^.. Given that all rhizomorphic lycopsids (sensu^20–23^) form a monophyletic group, and that extinct stigmarian rootlets were interpreted as modified leaves this suggested that the rootlets of all rhizomorphic lycopsids were modified leaves, including the rootlets of extant *Isoetes*^3^. The interpretation that the rootlets of extant *Isoetes* species were modified leaves was strikingly at odds with all previous descriptions of *Isoetes* rootlets that had always been interpreted as roots similar to the roots of other extant lycopsids^11,24–32^.

New evidence that is inconsistent with the modified shoot hypothesis has been reported since the seminal paper by Rothwell and Erwin^3^. First, the modified shoot hypothesis posits that the ancestral embryo condition in the rhizomorphic lycopsids lacked an embryonic root, but instead developed a single shoot axis that divided to give a typical shoot and modified rooting shoot axis that developed modified leaves (rootlets). However, embryo development in the early diverging rhizomorphic lycopsid, *Oxroadia* developed an embryonic root^20^. Therefore, the embryo of *Oxroadia* does not support the hypothesis that a branching event in the embryo produced a rooting shoot axis (rhizomorph) that developed root-like leaves (rootlets). Second, while the leaves of all plants species develop exogenously^33^, in a process that includes the outer-most layers of the shoot, roots of extant *Isoetes* originate endogenously^34^. Therefore, the endogenous development of rootlets is inconsistent with their interpretation as modified leaves^34^. Third, the discovery of the development of root hairs on rootlets of extinct rhizomorphic lycopsids that are identical to the root hairs that develop on extant lycopsids suggest that rootlets are root-like^2^. Together these three studies present an emerging body of evidence that is incompatible with the modified shoot hypothesis.

To independently test the modified shoot hypothesis for the origin of lycopsid roots, we evaluated gene expression data of the extant rhizomorphic lycopsid, *Isoetes echinospora*. We generated, to our knowledge, the first organ specific transcriptome of an *Isoetes* species incorporating RNA from the three main organs of the sporophyte: rootlets, leaves and corms. If *I. echinospora* rootlets are modified leaves as predicted by the modified-shoot hypothesis we would expect gene expression profiles to be similar in rootlets and leaves. If, on the other hand, *I. echinospora* rootlets are true roots as predicted by the lycopsid root hypothesis we would expect that gene expression profiles would be different between leaves and rootlets, and gene expression profiles would be similar between *I. echinospora* rootlets and roots of *Selaginella* species.

## Results

### Development of a protocol to propagate *Isoetes echinospora* in axenic culture

To define gene expression signatures in the organs of *I. echinospora*, a population of plants was collected from the wild (Fig. 1A) and protocols to grow the plants in axenic culture were developed. The collected plants were grown in the green house and male spores (microspores) and female spores (megaspores) were produced and then isolated. Megaspores and microspores were surface sterilised and germinated together in sterile liquid media to generate a population of sporophytes in axenic culture (Fig. 1B). Sporophytes were transferred to solid media three months after germination (Fig. 1C). A population of c. 50 *I. echinospora* plants were grown for approximately four months to a stage where plants were large enough to extract RNA from the three major organs; leaves, corm and rootlets (Fig. 2A).

**Fig. 1.**
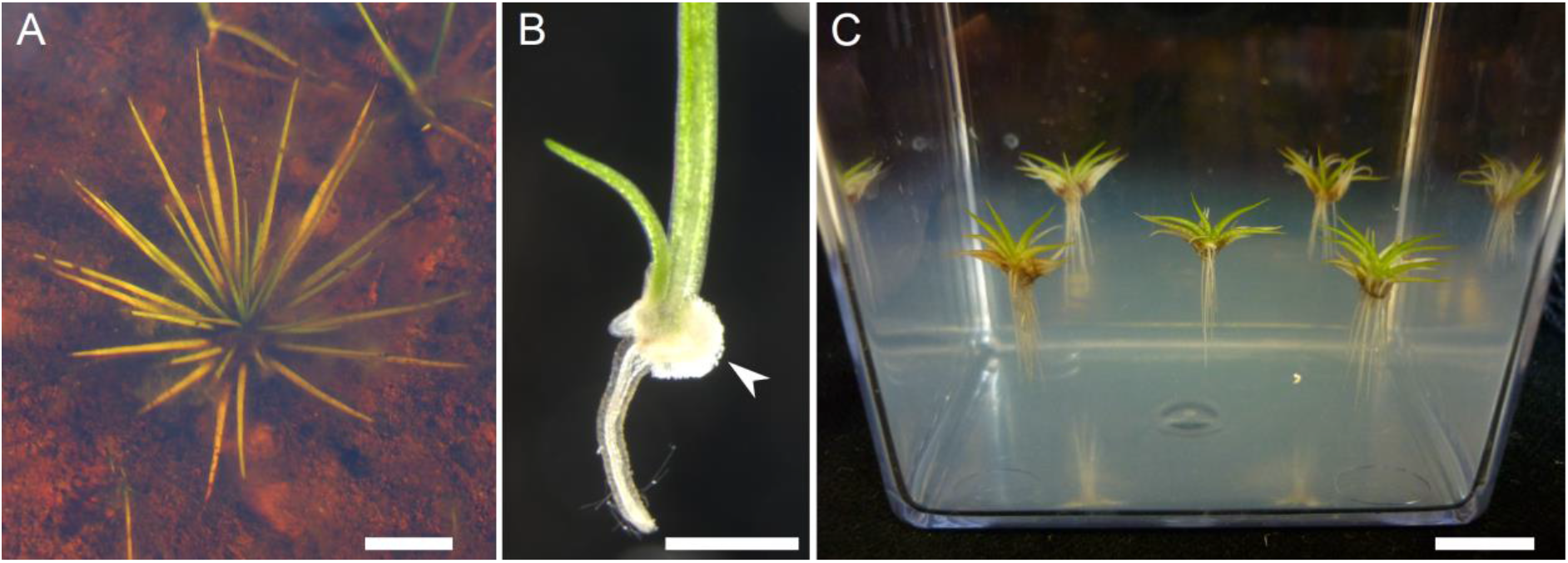
Growth of *I. echinospora* in axenic culture for RNA extraction. (*A*) *I. echinospora* growing submerged in its natural habitat, North West Sutherland (Scotland, UK). (*B*) Developing sporophyte emerging from the megaspore (highlighted with arrowhead) at the two leaf-, two rootlet-stage (*C*) *I. echinospora* sporophytes growing submerged on transparent solid media. Scale bars: A, approximately 2 cm; B, 1 mm; C, 1 cm.

**Fig. 2.**
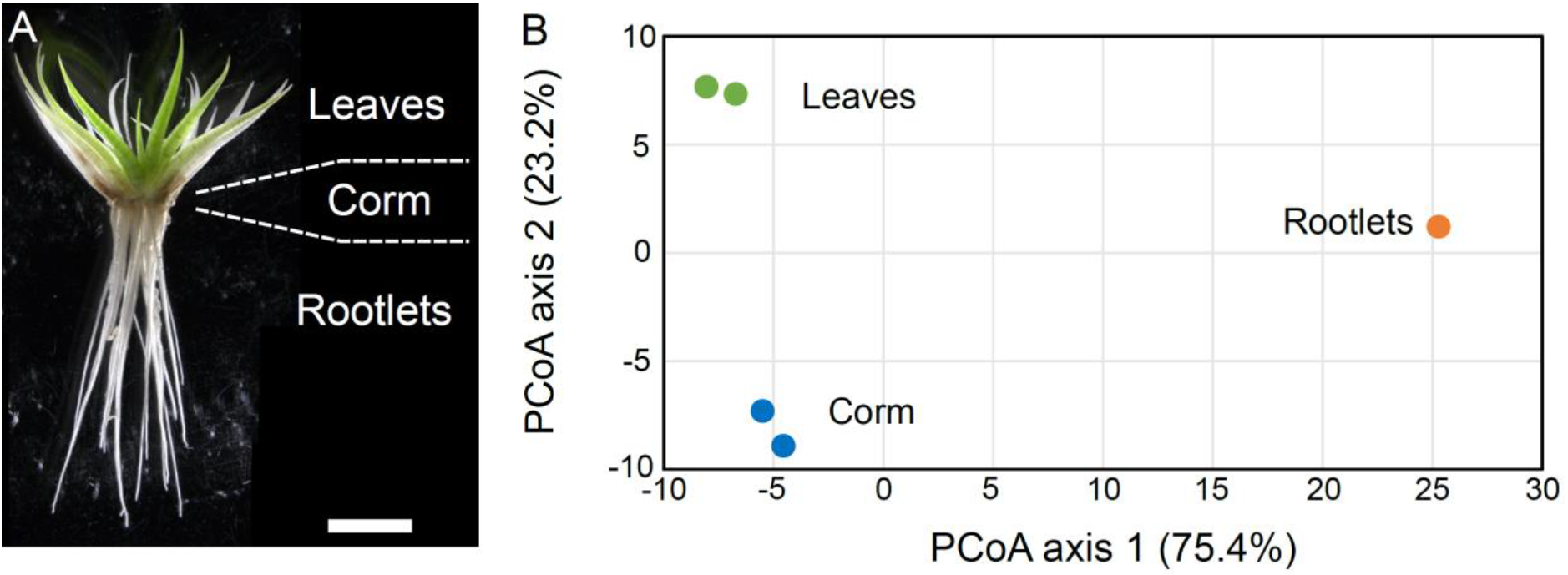
Gene expression profiles of leaves and rootlets are different in *I. echinospora*. (A) *I. echinospora* sporophyte at the stage when RNA was extracted from rootlets, corms and leaves. Scale bar 5 mm. Comparison of gene expression profiles by principle coordinate analysis (PCoA) in the transcriptome of *I. echinospora*. Two technical replicates of gene expression profiles of the corm and leaves and single replicate of rootlets of *I. echinospora*. Two leaf replicates, green. Two corm replicates, blue. Single root replicate, orange. Values on PCoA axes are shown in thousands. Values in brackets on each axis describe the percentage of total variance accounted for by each axis.

### Assembly of an *Isoetes echinospora* sporophyte transcriptome

A sporophyte transcriptome was generated for rootlets corms and leaves. RNA was isolated from each organ and sequenced. It was difficult to extract sufficient RNA from these plants because of the challenge in isolating viable spores, getting the spores to germinate, effecting fertilisation and getting sporophytes to develop in axenic culture. However, we extracted 1 technical replicate of rootlets and 2 technical replicates of corm and leaves. The raw reads for all samples were pooled, quality checked and assembled into contiguous transcripts. The assembled transcriptome comprised 113,464 transcripts with a mean sequence length of 940 base pairs (bp). There were 35,564 sequences over 1 kilobases (Kb) in the assembly, with an N50 of 1313 bp. Proteins were successfully predicted for c. 95% of the transcripts. To investigate the completeness our transcriptome we next performed a BUSCO^35^ analysis to investigate the number of conserved BUSCO^35^ groups in our transcriptome. BUSCO^35^ groups are near-universal single-copy orthologs. Identifing the percentage of BUSCO^35^ groups present in our *de-novo* transcriptome therefore provides a metric for the completeness of our transcriptome. Of the 430 total BUSCO^35^ groups searched for in the Viridiplantae dataset^35^, 318 (74.0%) were found complete, 87 (20.2%) were found fragmented and only 25 (5.8%) were missing. These metrics indicate that the transcriptome assembly was high quality. We next mapped the reads extracted from each of the three different organs; leaves, corms, and rootlets to calculate the abundance levels for each transcript in each of the three organs (Supplementary Table S1).

### Gene expression profiles are significantly different in *Isoetes echinospora* rootlets and leaves

If *I. echinospora* rootlets were modified leaves, as predicted by the modified shoot hypothesis, we might expect gene expression signatures to be similar in the rootlets and leaves. To test this hypothesis, we compared gene expression in rootlets, leaves and corms using a principal coordinate analysis (PCoA). The two leaf replicates, two corm replicates and the single rootlet sample were plotted on the first two PCoA axes (which together account for 98.6% of the variance in the sample (Fig. 2B)). The three tissue types are clearly distinct and separated in gene expression space. The first PCoA axis accounts for 75.4% of the variance in gene expression and it distinguishes leaves and corms from rootlets (Fig. 2B). The second PCoA axis accounts for 23.2% of the variance in gene expression and distinguishes all three tissues from each other (Fig. 2B). The PCoA indicated that gene expression profiles of rootlets and leaves are distinct and does not support the hypothesis that *I. echinospora* rootlets are modified leaves.

### Gene expression profiles of *Isoetes* rootlets clusters with gene expression of *Selaginella* and *Arabidopsis* roots

If the rootlets of *I. echinospora* are true roots we expected similarities in gene expression between rootlets and true roots of other land plant species such as the lycophytes *Selaginella moellendorffii* and the seed plant *Arabidopsis thaliana*. To compare gene expression between these species we first defined orthologous relationships between the genes of the three species using the OrthoFinder software^36,37^. This analysis identified 1,737 single copy orthologs in common between these species. Using these 1,737 orthologs we compared gene expression between the different species. We compared average gene expression between *I. echinospora* rootlets and leaves (this study) with the published gene expression in roots and leaves of *S. moellendorffii* ^38^ and roots and “aerial parts” of *Arabidopsis thaliana* (based on EMBL-EBI accession E-GEOD-53197). To compare gene expression between these different species and organs we subjected the gene expression dataset to a PCoA. The first three principal coordinates accounted for 95.7% of the variance in the dataset. Axis 1 accounted for 43.6% of the variance and separated the samples by species (Fig. 3A, B). Axis 2, accounted for 35.9% of the variance and distinguished the two lycophyte transcriptomes (*I. echinospora* and *S. moellendorffii*) from that of the seed plant *A. thaliana* (Fig. 3A, C). PCoA axes one and two therefore indicate that the majority of the differences in gene expression is accounted for by differences between species rather than between roots and leaves. PCoA axis 3 accounted for 16.2% of the variance and distinguished between leaves and roots in all species (Fig. 3B, C). Leaf samples clustered in the positive values and root samples clustered in the negative values of PCoA axis 3 (Fig. 3B, C). The clustering of the *I. echinospora* rootlet sample with both the roots of *S. moellendorffii* and *A. thaliana* on axis 3 (Fig. 3B, C) indicates that the gene expression signature of the rootlets of *I. echinospora* is similar to the the gene expression signature of both *S. moellendorffii* and *A. thaliana*. These gene expression data are consistent with the hypothesis that rootlets of *I. echinospora* are roots.

**Fig. 3.**
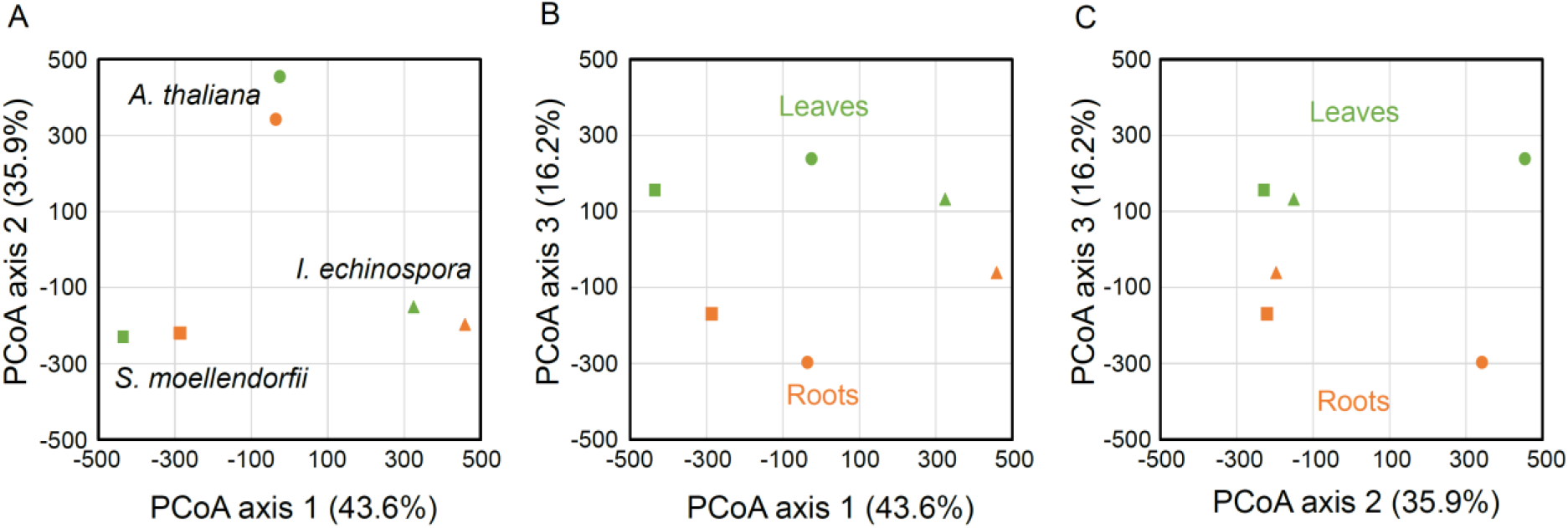
Comparison of root and leaf transcriptomes of *Arabidopsis thaliana, I. echinospora* and *Selaginella moellendorffii*. Comparison of gene expression profiles by PCoA in the transcriptomes of *A. thaliana*, circles, *S. moellendorffii*, squares and *I. echinospora*, triangles. Leaf samples coloured green, root samples orange. A, comparison of principal coordinate axis 1 and 2. B, comparison of principal coordinates axis 1 and 3. C, comparison of principal coordinate axis 2 and 3. Values in brackets on each axis describe the percentage of total variance accounted for by each axis. Axis 1 separates gene expression in the three species. Axis 2 distinguishes gene expression between the two lycophytes transcriptomes, *I. echinospora* and *S. moellendorffii* from *A. thaliana*. Axis 3 distinguishes between the leaf samples and the root samples in each transcriptome.

### The *RSL* root cell differentiation genes are expressed in *Isoetes echinospora* rootlets

To verify our findings that gene expression of *I. echinospora* rootlets were similar to those of the true roots of *S. moellendorffii* and *A. thaliana* we next determined the expression of the root-specific *ROOT HAIR DEFECTIVE SIX-LIKE* (*RSL*) genes in *I. echinospora. ROOT HAIR DEFECTIVE SIX-LIKE* (*RSL*) genes positively regulate the development of root hairs in euphyllophytes including *A. thaliana*^39–42^ and are expressed in *S. moellendorffii* roots^43,44^. *RSL* genes are markers for vascular plant roots because they are expressed at a much higher level in roots of *A. thaliana* (EMBL-EBI accession E-GEOD-53197) and *S. moellendorffii*^45^ than in leaves and shoots (Supplementary Table S1). We searched the *I. echinospora* transcriptome for RSL genes using the BLAST algorithm with RSL-specific queries. RSL sequences were identified in the *I. echinospora* transcriptome. A gene tree was generated and defined four *I. echinospora* RSL genes in two monophyletic groups (Fig. 4). There were three transcripts in the RSL Class I clade (106204; 101034; 092963) and a single transcript in the RSL Class II clade (095243). Average expression of the four RSL genes in rootlets was 4.24 transcripts per million (TPM) (Fig. 4). The average root expression was 5.78 TPM for the six RSL genes of *A. thaliana* (EMBL-EBI accession E-GEOD-53197), demonstrating similarities in expression of RSL genes between in *I. echinospora* and *A. thaliana*. In *I. echinospora*, expression of each RSL Class I transcript was higher in rootlets than in leaves (Fig. 4). Furthermore, the single *I. echinospora* Class II RSL gene transcript (095243) was expressed in rootlets and no expression was detected in the corm or leaves. These data indicate that RSL genes are preferentially expressed in the *I. echinospora* rootlets and not in leaf tissue, as in they are in *S. moellendorffii* of *A. thaliana* (Supplementary Table S1). These data are consistent with the hypothesis that *I. echinospora* rootlets are roots.

**Fig. 4.**
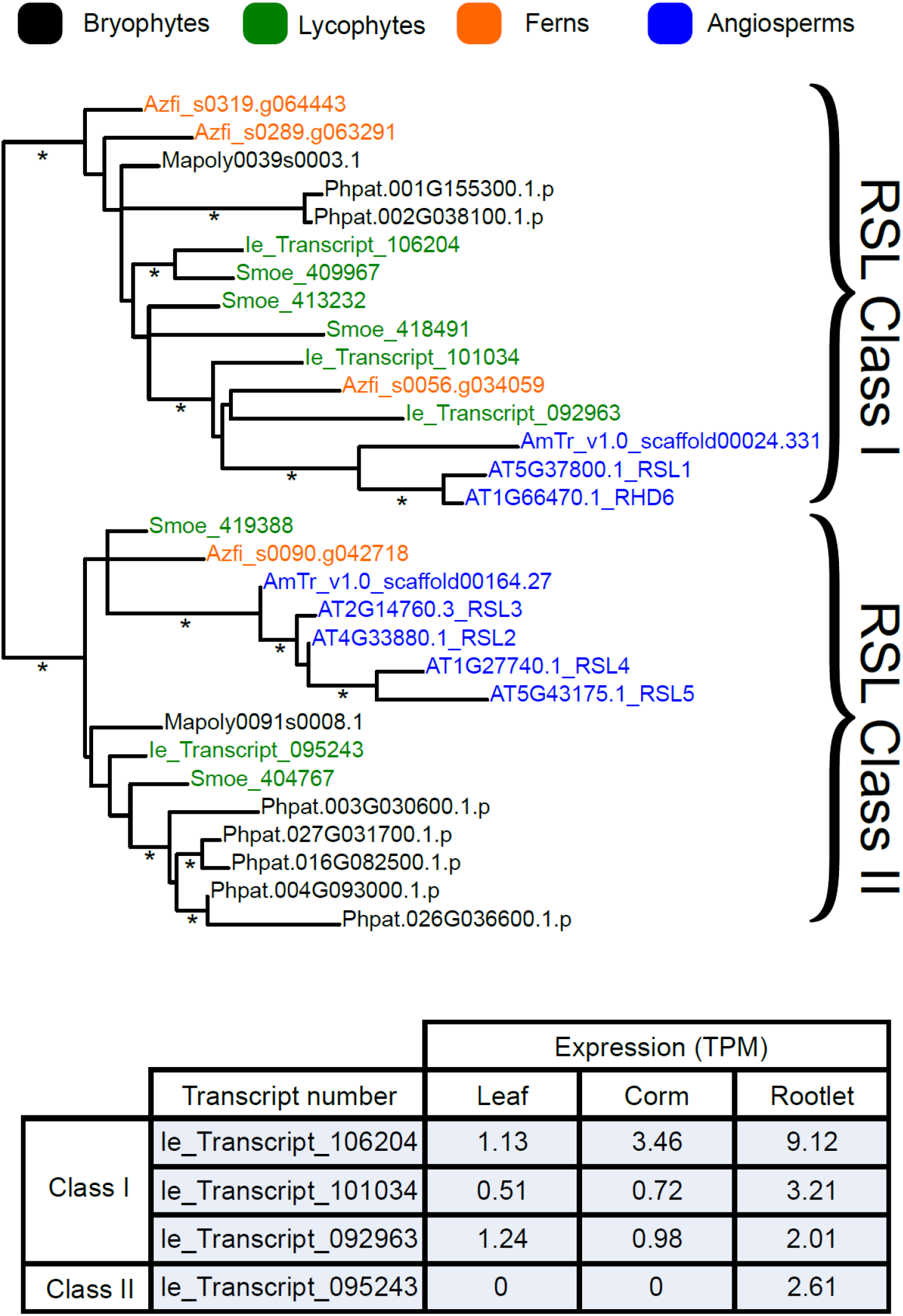
Gene tree analysis and expression analysis of *I. echinospora* RSL genes. Top, maximum likelihood gene tree of RSL genes generated in PhyML 3.0^87^. Gene names: black, bryophytes; green, lycopsids; orange, ferns; blue, angiosperms. The RSL genes are grouped into two monophyletic classes; Class I and Class II. Bottom, gene expression in the three RSL Class I genes and single RSL Class II gene in the *I. echinospora* transcriptome. Expression in transcripts per million (TPM) given for the single replicate of rootlets and as an average of the two technical replicates of leaf and corm. Species name abbreviations: Phpat, *Physcomitrella patens*; Mapoly, *Marchantia polymorpha*; Some, *Selaginella moellendorffii*; Sk, *Selaginella kraussiana*; Ie_Transcript, *Isoetes echinospora*; Sacu, *Salvinia cucullata*; Azfi, *Azolla filiculoides*; AmTr, *Amborella trichopoda*; AT *Arabidopsis thaliana*. * Indicate branches with over 0.85 aLRT SH-like support.

To verify that RSL genes are markers of vascular plant roots we investigated the *RSL* genes in *Azolla filiculoides*, a fern that develops roots with root hairs^30,46^, and *Salvinia cucullata* a fern that has secondarily lost roots with root hairs and instead modified leaves perform rooting functions^30,47,48^. We searched the *S. cucullata* and *A. filiculoides* genomes and proteomes^49^ for RSL genes using the BLAST algorithm with RSL-specific queries. A gene tree was constructed with the retrieved sequences and allowed us to identify 3 RSL Class I genes and a single RSL Class II gene in the *A. filiculoides* genome (Fig. 4, Supplementary Fig. S1). Consistant with their role in root development in *A. filiculoides* the RSL genes were expressed in the roots^46^. However, there were no RSL genes in the *S. cucullata* genome. *S. cucullate* sequences were identified in closely related basic-helix-loop-helix transcription factor subfamily XI^50,51^ but none were identified in RSL clade (Fig. 4, Supplementary Fig. S1). We conclude that the loss of RSL genes accompanied the evolutionary loss of roots with root hairs in *Salvinia cucullate*, which is consistent with RSL genes being markers of vascular plant roots. Furthermore, if the rootlets of *I. echinospora* were modified leaves, similar to the root-like modified leaves of *S. cucullate*, we might have expected that RSL genes would have also been lost from the *I. echinospora* genome. Instead, RSL genes are preferentially expressed in *I. echinospora* rootlets just as they are in *S. moellendorffii* roots. These data are consistent with the hypothesis that the *I. echinospora* rootlet is a root and not a modified leaf.

Taken together these data – the distinct gene expression profiles of the rootlets and leaves of *I. echinospora*, the similarity in expression profiles of orthologous gene preferentially expressed in rootlets of *I. echinospora* and roots of *S. moellendorffii* and *A. thaliana*, and the expression of the *RSL* genes in the rootlets of *I. echinospora* and roots of *S. moellendorffii* and *A. thaliana* – support the lycopsid root hypothesis which posits that *Isoetes* rootlets are roots and not modified leaves.

## Discussion

The homology of the rootlets of both extinct and extant rhizomorphic lycopsids have been contentious for the past 150 years, with two competing hypotheses. The first, interprets the rootlets as true roots similar to the roots of other lycopsids. The second, interprets rootlets as modified leaves. Despite the second hypothesis that posits that rootlets are modified leaves being widely accepted over the past 30 years^1,3,52^ there is a growing body of evidence^2,34^ that suggests that rootlets should be interpreted as true roots. Here we report the *de novo* transcriptome of *I. echinospora* that we used to test predictions of the two competing hypotheses. We discovered that expression profiles in *I. echinospora* rootlets and leaves were different. We showed that gene expression profiles of *I. echinospora* rootlets and *S. moellendorffii* and *A. thaliana* roots were similar. Finally, RSL genes involved in root cell differentiation are preferentially expressed in *I. echinospora* rootlets as they are in *S. moellendorffii* roots and the roots of euphyllophytes (A. thaliana, *Oryza sativa* and *Brachypodium distachyon*^39–42^). Taking these three pieces of evidence together, we conclude that *Isoetes* rootlets are true roots, like those of extinct and extant lycopsids and not modified leaves.

The new evidence presented here adds to the growing and extensive list of similarities between the rootlets of rhizomorphic lycopsids – *Isoetes* species and extinct taxa such as *Stigmaria* – and the roots of other lycopsids^2,20,34^. This growing body of evidence supports the hypothesis that rootlets are roots and not modified leaves. The rootlets of the rhizomorphic lycopsids and roots of all extant lycopsids are indeterminate radially symmetric axes that branch by isotomous dichotomy, develop endogenously within specialised structures, develop a root meristem with root cap and produce root hairs^31,32,34,53^. If the modified shoot hypothesis were correct it would have required the direct modification of a determinate leaf that did not branch, developed exogenously and was characterised by a ligule, stomata and dorsiventral symmetry into a rootlet. Each of these leaf characters would have had to be lost and all of the rootlet characters, which are shared among the lycopsids, would have had to evolve independently. By contrast, if the lycopsid root hypothesis is correct and rootlets are roots then there is no requirement for this large suite of character state changes. Instead, the only character transitions required to account for rootlet character states were the collateral positioning of the phloem, the regular rhizotaxy and rootlet abscission^4^. Although these three characters (collateral positioning of the phloem, the regular rhizotaxy and rootlet abscission) are predominately leaf characters, they are not exclusive to leaves; each has been described in the roots of other species of land plants. The collateral position of the phloem is found in Lycopodium roots including, *Lycopodium lucidulum, Lycopodium clavalum, Lycopodium obscurum* and *Lycopodium complanalum*^54^, regular rhizotaxy develops in *Ceratoptertis thalictroides, Cucurbita maxima* and *Pontederia cordata*^55^–^57^ and roots abscise in *Oxalis esculenta, Abies balsamea, Pinus strobus, Tsuga canadensis* and *Azolla* species^58–63^. Based on character transitions alone we suggest that the hypothesis that rootlets of the rhizomorphic lycopsids are roots, similar to other lycopsid roots, is a more parsimonious hypothesis than interpreting rootlets as modified leaves.

Our new evidence from the transcriptome of *I. echinospora* adds to the numerous traits that are common between the rootlets of rhizomorphic lycopsids and the roots of other lycopsids. It is not possible to rule out the hypothesis that all of these similarities in antomy, develop and now gene expression may be the product of convergent evolution. However, we suggest that it is more parsimonious to interpret the rootlets of the rhizomophic lycopsids as true roots than modified leaves.

The gene expression data from the *de novo I. echinospora* transcriptome are consistent with the hypothesis that the rootlets of the rhizomorphic lycopsids are roots and not modified leaves. We therefore interpret the rootlets of the rhizomophic lycopsids as roots developing from a unique root bearing organ; the rhizomorph^21,53,64^. This conclusion suggests that the dichotomously branching rooting axis is conserved among all lycopsids and a distinguishing character of the group. The dichotomous branching of these rooting axes has been conserved for over 400 million years and our comparative transcriptomic analysis suggest that the RSL genes function during root development in *Selaginella* and *Isoetes* has been conserved since these species shared a common ancestor at least 375 million years ago^65^. Our comparative analysis of the transcriptomes of extant lycophytes support the hypothesis that the rooting systems of extant *Isoetes* species and their extinct giant ancestors are homologous. The data also suggest that the development of the large rooting systems of the lycopsid trees that were an important component of the Palaeozoic flora and played a key role in changing the Earth’s Carbon Cycle were controlled by the same genes that regulate root development in their extant herbaceous descendants.

## Materials and Methods

### Plant collection and growth

Mature *I. echinospora* plants were collected from Loch Aisir and Loch Dubhaird Mor in September 2013 and 2014 from North West Sutherland (Scotland, UK) with the permission of the John Muir Trust and the Scourie Estate. *I. echinospora* plants were identified on the basis of their echinate megaspore ornamentation^66^. Mature *I. echinospora* plants were grown submerged in aquaria in Levington M2 compost topped with coarse gravel in a glasshouse at Oxford University at 18°C under a 16 h light: 8 h dark photoperiod.

### Growth of *I. echinospora* in axenic culture

RNA was extracted from plants grown in axenic culture to ensure that there was no RNA contamination from other organisms. A procedure was developed to surface sterilise *I. echinospora* spores and germinate a population of axenically grown plants, based on previously developed procedures^67–69^. Sporophylls were removed from the mature plant population growing in aquaria in September (2013 and 2014) when sporangia were mature^67^. Using forceps (under a Leica M165 FC stereo microscope) mega-and micro-sporangia were isolated from sporophylls. Intact sporangia were washed in 1% (v/v) sodium dichloroisocyanurate (NaDCC) for 5 min. Sporangia were broken and loose spores were washed in 0.1% NaDCC for a further 5 min. Following the NaDCC washes, loose spores were rinsed for 5 min three times in ddH2O. Microspores were centrifuged for 5 min at 5000 rpm between washes). Once sterilised, mega and micro-spores were mixed together in ddH2O in a Petri dish. Petri dishes were sealed with parafilm, and incubated in darkness at 4°C for 2 wk. After 2 wk, Petri dishes were moved to a 16 h light: 8 h dark photoperiod at 18°C. Approximately 30% of surface sterilised megasporangia contained megaspores that germinated, and within these megasporangia c. 25% of the total megaspore population germinated. It was possible to identify germinating megaspores because cracking of the megaspore wall was visible and the presence of archegonia on the megagametophyte. Once fertilisation occurred, developing sporophytes were identified by the presence of the first leaf. Sporophytes were left to continue to grow in ddH2O water until the two leaf two rootlet stage when they were moved to magenta boxes containing; ½ Gamborg’s medium^70^, supplemented with 1% phytogel (Sigma). Plants were embedded in Gamborg media and submerged in liquid Bold’s Basal Medium (Sigma, UK).

### RNA extraction and sequencing

Total RNA was extracted from root, corm and leaf tissues from c. 50 *I. echinospora* plants. Total RNA from leaves (two independent replicates), corm (two independent replicates) and rootlets (one replicate) was extracted with the RNeasy plant mini kit (Qiagen). On-column DNase I treatment was performed with RNase-free DNase I (Qiagen), according to the manufacturer’s instructions. cDNA was synthesised with ProtoScript II reverse transcriptase (New England Biolabs) according to the manufacturer’s instructions, using oligo(dT) primer. Total cDNA samples were quantified with a Nanodrop ND-1000 spectrophotometer. RNA purity and quality were checked with an Agilent 2100 Bioanalyzer. cDNA was sequenced by the High-Throughput Genomics Group at the Wellcome Trust Centre for Human Genetics, University of Oxford. Sequencing resulted in 195,072,304 paired end reads separated into five samples: 2 leaves samples (35,718,157; 35,555,048 paired end reads), 2 corm samples (38,728,989; 44,379,751 paired end reads) and one rootlet sample (40,690,359 paired end reads). The raw read libraries have been deposited under SRP135936 on the NCBI Sequence Read Archive.

### *De novo* transcriptome assembly, protein predictions and expression analysis

Raw reads were quality trimmed using Trimmomatic-0.32^71^, to remove remaining Illumina adaptors and low quality tails. Ribosomal RNA was filtered out using Sortmerna-1.9^72^ and error corrected using BayesHammer (SPAdes-16 3.5.0)^73^ (with setting --only-error-correction) and Allpaths-LG-4832^74^ (with setting PAIRED_SEP=-20 and ploidy = 2). Reads were normalised using Khmer-0.7.1 with a khmer size of 21. Before assembly, paired end reads were stitched together using Allpaths-LG-4832^74^. A *de novo* transcriptome assembly was made with the cleaned, stitched reads using SGA^75^, SSPACE-v3^76^, and CAP3^77^. Finally assembled scaffolds were corrected using Pilon-1.6^78^. The Transcriptome Shotgun Assembly project has been deposited at DDBJ/EMBL/GenBank under the accession GGKY00000000. The version described in this paper is the first version, GGKY01000000. Proteins were predicted from the *de novo* transcriptome assembly using GeneMarkS-T^79^, Prodigal^80^ and Transdecoder (part of the Trinity assembly program^81^), proteins were deposited on Zenodo (http://doi.org/10.5281/zenodo.3574570). A BUSCO analysis^35^, using BUSCO 3.1.0 and the *viridiplantae_odb10* database.

### Comparison of gene expression between *I. echinospora* organs

Using the sporophyte transcriptome assembly we next mapped the reads from the three organ libraries – leaves, corms, and rootlets – to the transcriptome to measure the expression levels of each transcript in the three tissues using Salmon^82^. To investigate the similarities between gene expression in the different organs we carried out a PCoA on the three organ types. Euclidean distances were derived from the expression of all transcripts (TPM) in each organ and were subjected to PCoA in PAST^83^ using a transformation exponent of 2.

### OrthoFinder analysis and comparison of gene expression between *I. echinospora, S. moellendorffii* and *Arabidopsis thaliana*

Orthologous relationships between *I. echinospora, S. moellendorffii* and *A. thaliana* proteins were determined using OrthoFinder^36,37^. OrthoFinder was run with *I. echinospora* proteins and protein datasets for 57 species from Phytozome (full list of species in Supplementary Table S2) including the Rhodophyta *Porphyra umbilicalis*, seven species of chlorophytes, the bryophytes *Marchantia polymorpha* and *Physcomitrella patens*, the lycophytes *Selaginella moellendorffii* and 46 angiosperm species. This analysis resulted in the identification of 38,217 orthogroups, accounting for 82.6% of all genes included in the analysis (The results of the OrthoFinder analysis were we deposited on Zenodo, http://doi.org/10.5281/zenodo.3574570).

To compare gene expression between *I. echinospora, S. moellendorffii* and *A. thaliana* we identified single copy orthologs between these species based on the OrthoFinder^36,37^ analysis. In total, 1,737 single copy orthologs were found between the three species. Using these 1,737 orthologs we contrasted gene expression between the different species. We investigated average genes expression between *I. echinospora* rootlets and leaves (this study) with the published average gene expression between roots and leaves of *S. moellendorffii*^38^ and *A. thaliana. A. thaliana* gene expression was based on average gene expression in “aerial part” and “root” of 17 different natural accessions (EMBL-EBI accession E-GEOD-53197). To investigate similarities in gene expression between these 1,737 orthologs we carried out a PCoA in PAST^83^. Euclidean distances were derived from the Log10 transformed gene expression of the 1,737 orthologs (Supplementary Table S1). Euclidean distances were subjected to a PCoA in PAST^83^, using a transformation exponent of 4.

### Phylogenetic Analyses

Phylogenetic analyses were carried out on the RSL genes. Blast queries were assembled based on previously published gene trees of RSL genes^50^. Sequences were used to blast the protein databases of the; *Marchantia polymorpha* “primary” (proteins) (version 3.1, November, 2015), *Physcomitrella patens* “primary” (proteins) (version 3.0, January 12, 2014), *Selaginella moellendorffii* “primary” (proteins) (version 1.0, January 12, 2014), *Amborella trichopoda* (proteins) (version 1.0, 2013) and *Arabidopsis thaliana* “primary” (proteins) (TAIR10) on the http://marchantia.info/blast/server. Two fern protein databases were also searched; *Azolla filiculoides* protein v1.1 and *Salvinia cucullata* proteins v1.2 ^49^ as well as the predicted proteins from the *I. echinospora* transcriptome generated in this study. All proteins were aligned in MAFFT^84,85^, manually edited in Bioedit^86^. Maximum likelihood gene trees were generated in PhyML 3.0^87^, using Jones, Taylor and Thorton (JTT) amino acid substitution model. To verify the absence of *RSL* genes in the *S. cucullate* the genomes and proteomes of *A. filiculoides* and *S. cucullate* were searched by Blast using the *A. thaliana* protein sequence RSL1 (AT5G37800) using an E-value cut off of 1E-15. A gene tree was generated as described above including the addition of *A. thaliana* protein sequence from subfamilies VIIIb and XI^50,51^ (Fasta alignments files for both gene trees we deposited on Zenodo, http://doi.org/10.5281/zenodo.3574570).

## Acknowledgements

This research was supported by a Biotechnology and Biological Sciences Research Council (Grant BB/J014427/1) Doctoral Training Partnership Scholarship and the George Grosvenor Freeman Fellowship by Examination in Sciences, Magdalen College (Oxford) to A.J.H. L.D. was funded by a European Research Council Advanced Grant (EVO500, contract 250284), European Commission Framework 7 Initial Training Network (PLANTORIGINS, project identifier 238640) and European Research Council Grant (De NOVO-P, contract 787613). S.K. and D.E. were funded by a European Union’s Horizon 2020 research and innovation program under grant agreement number 637765 to

S.K. S.K. is a Royal Society University Research Fellow. We thank the High-Throughput Genomics Group at the Wellcome Trust Centre for Human Genetics (funded by Wellcome Trust grant reference 090532/Z/09/Z) for the generation of sequencing data. We are grateful to the John Muir Trust and the Scourie Estate for providing permission to collect *I. echinospora*, and to Dr Heather McHaffie (Royal Botanic Garden Edinburgh) for assistance identifying *I. echinospora*.

## Competing interests

The authors declare no competing interests.

## Data availability

The raw read libraries have been deposited under SRP135936 on the NCBI Sequence Read Archive. Transcriptome Shotgun Assembly project has been deposited at DDBJ/EMBL/GenBank under the accession GGKY00000000. The orthofinder analysis, predicted protein sequences in the I. echinopsora transcriptome, and fasta alignment files for gene trees were deposited at Zenodo, http://doi.org/10.5281/zenodo.3574570.

---

**Supplementary Figure S1.**
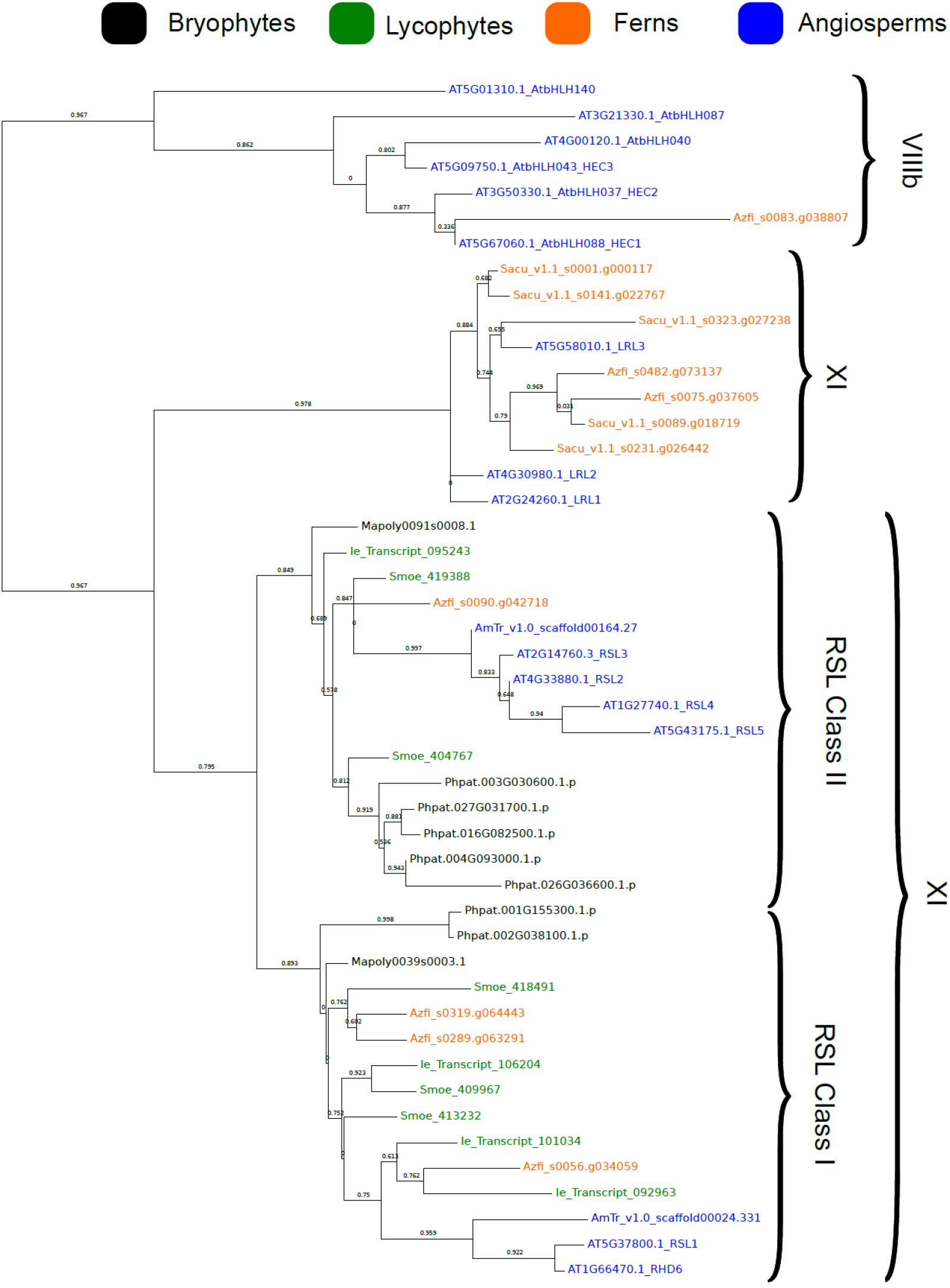
There are no RSL genes in the *Salvinia cucullata* genome or proteome. Gene tree analysis of RSL and related basic Helix-Loop-Helix (bHLH) transcription factors in a subset of land plant species. There are not *S. cucullata* genes in the RSL clade instead closely related *S. cucullate* genes are members of subfamily XI. Maximum likelihood gene tree of bHLH transcription factors generated in PhyML 3.0. Gene names: black, bryophytes; green, lycopsids; orange, ferns; blue, angiosperms. The RSL genes are grouped into two monophyletic classes; Class I and Class II. Species name abbreviations: Phpat, *Physcomitrella patens*; Mapoly, *Marchantia polymorpha*; Some, *Selaginella moellendorffii*; Sk, *Selaginella kraussiana*; Ie_Transcript, *Isoetes echinospora*; Sacu, *Salvinia cucullata*; Azfi, *Azolla filiculoides*; AmTr, *Amborella trichopoda*; AT *Arabidopsis thaliana*. Branch support as Shimodaira-Hasegawa-like approximate likelihood ratio tests.

## Notes

http://doi.org/10.5281/zenodo.3574570

https://www.ncbi.nlm.nih.gov/nuccore/GGKY00000000.1/

